# Intercellular positive-feedback loops promote the evolutionary stability of microbial cooperative behaviors

**DOI:** 10.1101/571562

**Authors:** Ishay Ben-Zion, Avigdor Eldar

## Abstract

Microbial cooperation enables groups of conspecific cells to perform tasks that cannot be performed efficiently by individual cells, such as utilization of various secreted ‘public-good’ molecules, communication via quorum-sensing, or the formation of multicellular structures. Cooperation is often costly and therefore susceptible to exploitation by ‘cheater’ cells, which enjoy the benefit of cooperation without investing in it. While population structure is key to the maintenance of cooperation, it remains unclear whether other mechanisms help in stabilizing microbial cooperation. Like other microbial traits, cooperation is often governed by complex regulatory networks, and one reoccurring motif is an ‘intercellular positive-feedback loop’, where a secreted molecule, e.g. a public-good or a quorum-sensing signaling molecule, activates its own production in all surrounding cells. Here we investigate the role of intercellular feedbacks in the maintenance of bacterial cooperation. We combine theory with a synthetic-biology approach, using swarming motility of *Bacillus subtilis* engineered variants, to compare the response of ‘open-loop’ and feedback cooperators to the presence of cheaters. We find that positive feedbacks on cooperative behaviors – either directly or through a feedback on quorum-sensing – maintain cooperation in a broader range of environments, relieving the requirement for a strong population structure. Our results directly demonstrate the stabilizing effect of intercellular positive feedbacks on cooperative behaviors, and suggests an explanation for their abundance in regulatory networks of bacterial cooperation.

## Introduction

Microbial behavior is guided by complex regulatory gene networks. These networks integrate information from the environment with information on the cell state in order to reach a “decision” on whether or not to express a certain trait and to which extent. Understanding the function of these regulatory networks and the design principlesiding them is a major focus of systems biology [1,2]. Specifically, feedback loops have attracted much attention and studied theoretically and experimentally. Negative feedbacks were shown to stabilize systems output [3,4], to reduce noise [5] or to speed up response [6], while positive feedbacks were mostly studied as modules that create bistability [7–9], or sharp irreversible transitions during differentiation [10,11].

The study of regulatory networks is also crucial for our understanding of the dynamics of microbial social behaviors and specifically cell-cell signaling and cooperation. Many microorganisms exhibit cooperative behaviors – the act of one cell is beneficial to the whole group of cells, but costly to the individual cell [12,13]. The canonical example is secretion of ‘public-good’ molecules, such as extra-cellular metabolic enzymes [14–16], surfactants [17–20] and virulence factors [21–23]. Many microbial cooperative behaviors are guided by quorum-sensing cell-cell communication systems, where cells regulate cooperative behaviors in a density-dependent manner through secretion and response to signaling molecules [24].

The dependence of cooperative behaviors on secreted molecules, either public goods or signaling molecules, allows for the formation of intercellular (group-wide) feedback loops, whereby the secretion rate of a cell is determined by the (local) external concentration of the secreted molecule. This creates an association between the individual investment in cooperation and the mean investment in its local environment [25]. Both positive and negative intercellular feedback loops are prevalent in the regulatory networks of microbial cooperative behaviors, and especially in the secretion of public-good molecules. First, production of many public-good exoenzymes is directly inhibited by their product (e.g., [26]), or regulated by the stress-response system [19]. Both of these forms of regulation will lead to the formation of an ‘intercellular negative-feedback’ loop. In the latter case, the negative feedback is formed by virtue of the reduction of stress levels through the cooperative act. Second, public-good molecules sometimes regulate the transcriptional activation of their producing genes, forming an ‘intercellular positive-feedback’ loop. Iron-scavenging siderophores provide one example for such a design, since the binding of ferri-siderophores to their specific receptors for reentering the cell also activates the siderophore synthesis pathway [27–30]. Another example for this positive-feedback design is the secretion of bacteriocins, which benefit all immune bacteria. Bacteriocin production is sometimes triggered by self-induced DNA damage [31–33], thereby forming an intercellular positive feedback on public goods. Finally, many quorum-sensing systems, in addition to their regulation of cooperative behaviors, activate the production of their own signaling molecules, and this is another form of an intercellular positive feedback on cooperation [34,35].

Intercellular feedbacks may lead to different effects than intracellular ones. For example, unlike intracellular positive feedbacks, intercellular positive feedbacks should decrease cell-to-cell variations and homogenize a population [36–38], yielding bistability in the level of the population that could be either all ‘ON’ or all ‘OFF’. Intercellular positive feedbacks were suggested to allow population synchronization [33,34], to speed-up QS response [39,40] and sharpen its density dependence [34,41], and to allow spatial wave propagation [42]. These suggested roles of intercellular positive feedbacks disregard the social conflict and the intrinsic susceptibility of microbial cooperative behaviors to exploitation and elimination by non-cooperative ‘cheater’ mutants – variants which do not invest in cooperation but enjoy its benefits. Such cheaters, if kept unchecked, will invade a cooperative population and lead to its demise. A key question in microbial ecology and evolution is therefore how cooperative traits are maintained in evolution. Theoretically, kin-selection has been a leading candidate for this evolutionary maintenance [12], and may very well fit the high level of population structure and relatedness observed in microorganisms [43]. However, there is an ongoing debate whether other mechanisms, such as association with private good [44], policing [45], or snow-drift interactions [16], can replace or add up to kin-selection in maintaining cooperation (reviewed in [12]). The effect of microbial regulatory-network structure on the evolutionary maintenance of cooperation has been less studied [35,46].

Here we use a combination of mathematical models and a synthetic biology approach to show that intercellular positive feedbacks stabilize cooperation by reducing cooperative investment around cheaters, as was proposed for QS regulation by simulations [46] and indirect experiments [35]. We present a model of feedback regulation that demonstrates that positive feedbacks extend the range of environments in which cooperation is evolutionarily stable, and this stabilizing effect positively depends on feedback strength. We construct and calibrate engineered variants of public-good regulation, with and without feedback, in *B. subtilis,* and show that the stabilizing effect of positive feedbacks is due to a positive association between the individual’s cooperative investment and the mean investment in its local environment. We further dissect the two advantages of this association; a reduced intra-group disadvantage, and an increased inter-group advantage, and use a simple structured population assay to validate the broadening of the range of environments where cooperation is stable.

## Results

### A simple mathematical model demonstrates that an intercellular positive feedback on public-goods cooperation increases stability against cheater mutants

To study the effect of intercellular feedbacks on the evolutionary stability of cooperative behaviors, we devised a simple mathematical model of feedback-regulated public goods (Fig. 1A). We assume that the public-good production rate *x* of an individual bacterium is linearly regulated by the local concentration of public goods (which in steady state is proportional to the average public-good production rate *x̄*, assuming constant cell density, see Fig. 1B and supplementary text for derivation):

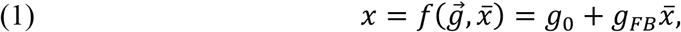

Where 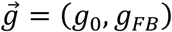 is a set of values that determine the form of the regulatory function *f*; *g*_0_ > 0 is the basal public-good production rate, and −1 < *g*_*FB*_ < 1 (positive for positive feedback and negative for negative feedback) is the feedback strength that measures the effect of the public good on its own production rate.

**Figure 1.**
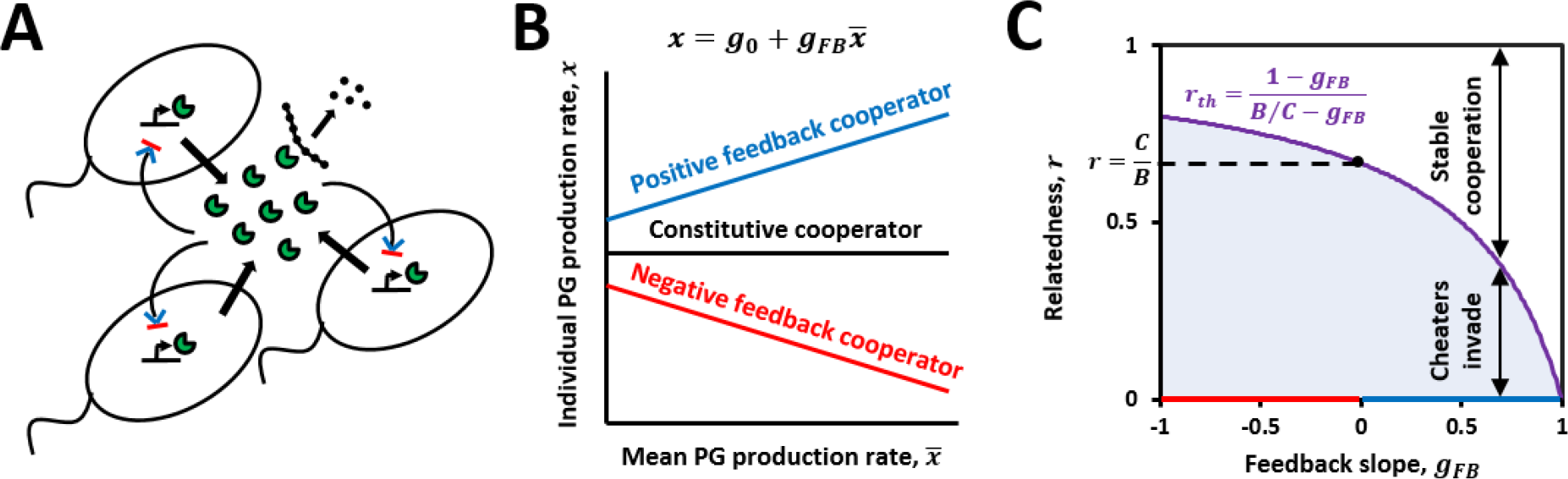
A mathematical model demonstrates that a positive-feedback structure improves evolutionary stability against cheaters. **(A)** The model describes bacterial secretion of public goods (PG) (here, exoenzymes that degrade complex nutrients into simpler nutrients that can enter the cell). Feedback control leads to either activation (blue arrow heads) or repression (red bar-heads) of PG production by their average concentration in the environment. **(B)** PG production rate, *x*, is assumed to be linearly dependent on external PG concentration (which at steady state is proportional to the mean production rate, *x̄*, see supplementary text). **(C)** Calculated threshold relatedness (*r*_*th*_, purple curve), for evolutionary stability against cheating, as a function of feedback slope, *g*_*FB*_ (blue/red regions for positive/negative feedbacks respectively). For every feedback slope, *g*_*FB*_, there is a threshold of relatedness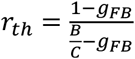 (purple line), above which cooperation is stable against cheater invasion and below it cheaters invade. Dashed line represents the Hamilton rule threshold for unregulated cooperation 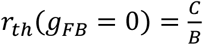. The benefit to cost ratio parameter used here is 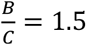

For simplicity, we assume a linear fitness function, where the fitness *W* of each individual decreases with its own public-good production rate (with a cost parameter *C*) and increases with the mean public-good production rate in its surroundings (with a benefit parameter *B*):

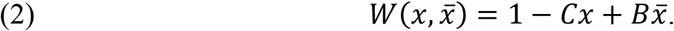

We tested the stability of cooperators with different regulatory strategies in the plane of (*g*_0_ × *g*_*FB*_) to invasion by public-good null cheater mutants, i.e. (*g*_0_ = 0, *g*_*FB*_ = 0). We assumed a simple population structure such that the frequency of invaders in a patch is either zero or equals the structural relatedness coefficient *r* [47–49]. We therefore compared the fitness of an invading cheater in a patch with a frequency *r* of cheaters to the fitness of a cooperator in a pure cooperator patch. We found that invasion success is independent of the basal public-good production rate *g*_0_ of the resident strategy, and depends only on the feedback strength *g*_*FB*_. The condition for stability of the strategy (*g*_0_, *g*_*FB*_) to invasion by cheaters is that the relatedness should be higher than a threshold that depends on *g*_*FB*_ (Fig. 1C):

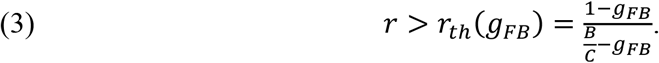

Figure 1C shows this threshold (purple line) as a function of the feedback strength *g*_*FB*_. The dashed line is the Hamilton rule threshold for constitutive (‘open-loop’) cooperation: 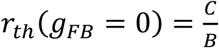. One can see that as *g*_*FB*_ becomes larger, the range of relatedness levels that allow cooperation to be stable against cheating becomes wider. Thus, a positive-feedback structure (*g*_*FB*_ > 0) promotes cooperation by weakening the demand of Hamilton’s rule for constitutive cooperation and allowing stable cooperation in a wider range of population structures, while a negative-feedback structure (*g*_*FB*_ < 0) makes this demand more stringent. We note that one can recapture Eq. (3) using the standard Hamilton rule, 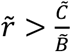, but *r̃*, *C̃*, *D̃* would be different than the structural relatedness *r* and our benefit and cost parameters *C*, *B* [50]. Interestingly, the same relation between feedback strength and the threshold relatedness is found when considering local evolutionary stability to small mutations (see Figs. S1,S2 and supplementary text).

Positive-feedback cooperators are better immune to cheating since they “sense” the presence of cheaters (when they sense a low concentration of public goods) and reduce their investment in cooperation with the increase in the frequency of cheaters in the population. Negative-feedback cooperators, on the other hand, invest even more in cooperation when there are cheaters around. The reduction in cooperative investment of positive-feedback cooperators helps them in two ways. First, they suffer a smaller cost when mixed with cheaters (and this cost decreases with their frequency in the group), reducing the intra-group disadvantage of cooperators. Second, cheaters in these mixed social groups, and especially in cheater-dominated groups, gain a smaller benefit, and this increases the inter-group advantage of cooperators.

How could these predictions be extended to feedbacks in QS regulation? In a QS design, where the binding of self-produced signals to their cognate receptors activates cooperative behaviors, a null mutation in the response genes yields a signal-blind cheater genotype. In many cases, QS response and signaling genes are co-transcribed as an operon [34], and therefore response-null mutants could also be signal-null mutants. In the supplementary text, we extend our model and analyze stability of two QS strategies; with and without a feedback on the QS signal, to invasion by response-null mutants and full QS-null mutants (response- and signal-null). We show that the simple QS design by itself provides an increased immunity against full QS-null mutants because QS-cooperators can "sense" their presence (by sensing low external signal concentrations) and reduce cooperative investment accordingly. However, simple QS-cooperators cannot "sense" the presence of response-null mutants that have a functional signaling gene. These mutants continue to produce QS signals and coerce cooperators to have full investment in cooperation. However, adding a feedback on the QS signal improves stability against these response-null mutants, and the resulting condition for stability is identical to the simpler case of auto-regulated public goods (Eq. (3)). Thus, the stabilizing effect of the intercellular positive-feedback structure applies here as well, improving the stability against response-null mutants that pose a threat to simple QS cooperation.

### Synthetically altered regulatory networks of QS in *Bacillus subtilis*, with and without feedback, can be calibrated to display comparable cooperation levels

To experimentally test whether positive feedbacks improve stability to cheating, we have synthetically manipulated the regulatory network design of the Surfactin production pathway of *B. subtilis*. Surfactin is a secreted surfactant lipo-peptide, which enables *B. subtilis* to perform swarming motility on semi-solid agar [51], among other roles [52–55]. Surfactin production is cooperative and can be exploited by non-secreting cheater mutants during swarming [17,18]. Surfactin is produced by an enzymatic complex that is coded by the *srfA* operon [56]. This operon is activated by the transcriptional regulator ComA [57], part of a two-component system (ComP-ComA). The transmembrane receptor ComP phosphorylates and activates ComA upon binding by the quorum sensing signal ComX [58]. This signal is a modified peptide which is produced from the *comX* gene and further cleaved and modified by the product of the *comQ* gene [57]. The *comQXP* genes are known to be co-transcribed as an operon, with constitutive expression [59] (see Fig. 2A for an illustration of the pathway).

**Figure 2.**
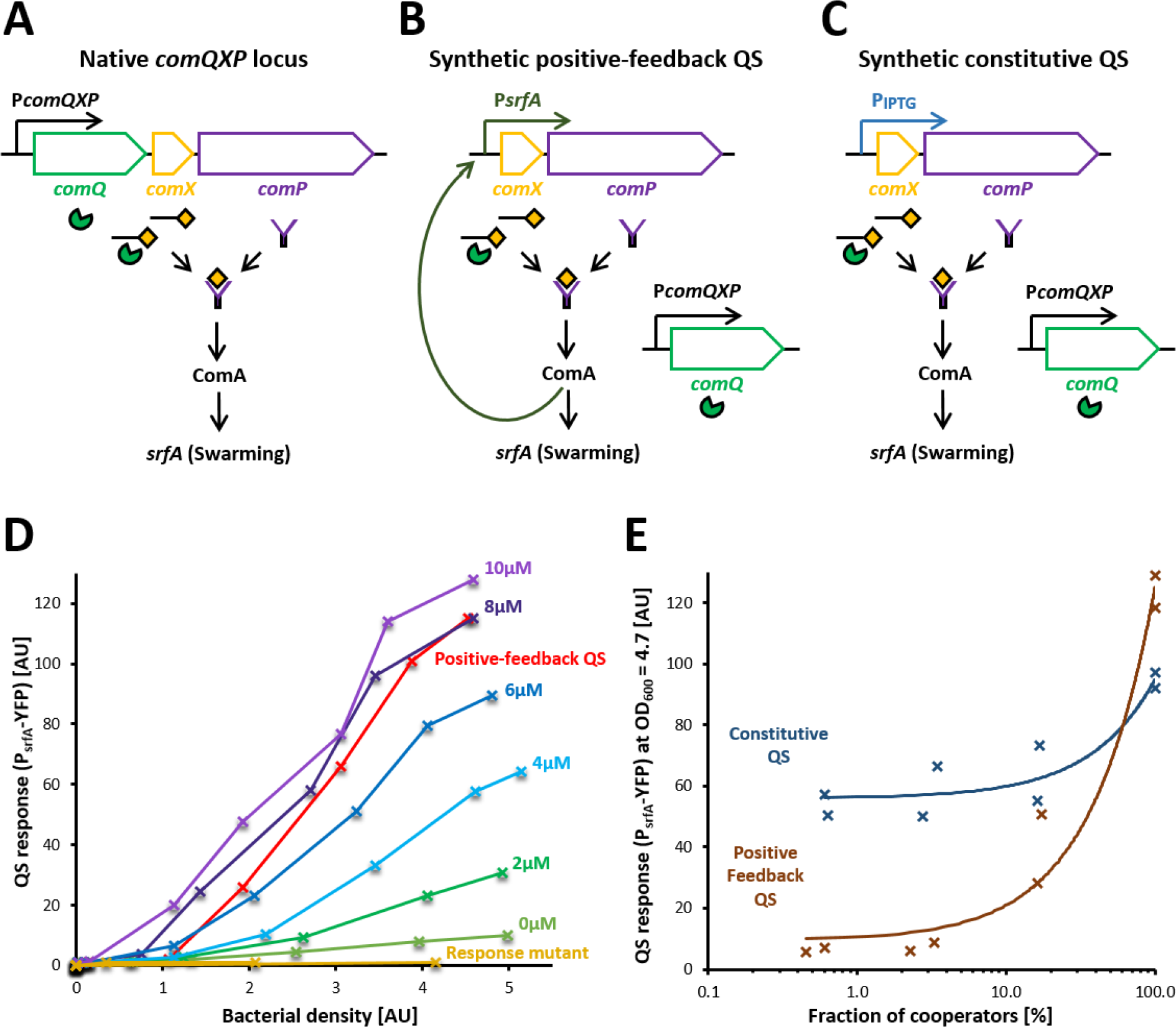
Design and gene expression of synthetic constitutive and positive-feedback QS systems. **(A-C)** Schemes of the native ComQXP QS system (A), the positive-feedback QS system (B, with *comXP* under a ComA-regulated promoter), and the inducible constitutive (‘open-loop’) QS system (C, with *comXP* under an IPTG-inducible promoter). **(D)** Density dependence of QS response for a response-null mutant (∆*comA*, AES3412, orange), positive-feedback-QS strain (AES3430, red) and an inducible constitutive-QS strain at multiple IPTG levels (AES3429, other colors, see legend). QS response is measured in liquid mono-cultures as the median of P_srfA_-YFP fluorescence values taken using flow cytometry (Methods) and displayed as a function of bacterial density (measured by counting flow-cytometer events, calibrated to OD_600_ values, Methods). One biological repeat is presented for each condition. **(E)** QS response of cooperators in liquid co-cultures with their response mutants. The QS response at a bacterial density corresponding to OD_600_ = 4.7 is presented for the constitutive QS-cooperator (AES3439, blue) and positive-feedback QS-cooperator (AES3440, brown) in co-cultures with their corresponding response-null mutants (AES3425 and AES3427, respectively). Data corresponds to two biological repeats of each strain. Lines are linear model fits to the data, with *R*^2^ = 0.85 and *R*^2^ = 0.97 for constitutive and positive-feedback QS-cooperators respectively. Note the logarithmic scale of the x-axis.

In order to compare the sensitivity of feedback and constitutive designs to cheating we modified the regulatory network of the comQXP system. To introduce a positive-feedback loop design, we placed the *comXP* part of the operon under the control of a copy of the *srfA* promoter (which is activated by ComA) (Fig. 2B). The native state of the system is constitutive; however, in order to properly calibrate the two systems we placed *comXP* under the control of an IPTG-inducible promoter (Fig. 2C). Each of these two constructs was then inserted into two backgrounds; into a *comXP* knockout strain to create cooperators, and into a *comXPA* knockout strain to create response-null mutants with functional signaling genes (*comQ* and *comX*). This setup allowed us to test the theoretical predictions that a positive-feedback loop on the QS signal improves the stability against response-null mutants.

Next, we calibrated the systems. To monitor cooperative secretion of surfactin, we introduced a genomic YFP transcriptional reporter of the *srfA* operon (P_srfA_-YFP) [60] into the various strains. By modifying IPTG concentration we were able to calibrate the system such that the two cooperators had similar cooperation levels (as measured by P_srfA_-YFP expression in shaken liquid cultures that do not require swarming motility, Methods). Fig. 2D describes the results of this calibration. Except for the response-null mutant, which showed no QS response, all other time-series displayed density dependence, as expected from QS response. For the constitutive QS strain, the density dependence slope increases with increasing IPTG levels, as higher bacterial density is needed to elicit QS response when signals and receptors are weakly expressed. The expression of the positive-feedback QS strain is comparable to the constitutive QS strain at an IPTG concentration between 6-8μM.

### Positive-feedback cooperators display a positive frequency-dependent cooperative investment

After choosing an IPTG concentration (7μM) that makes the two strains comparable in mono-cultures, we tested their cooperative investment level when each is co-cultured with its respective response-null mutant (genetically identical to cooperators except an additional *comA* knockout). Fig. 2E presents P_srfA_-YFP expression of cooperators in these co-cultures, estimated at a chosen bacterial density, corresponding to OD_600_ = 4.7 (Methods). As expected, the cooperative investment of the constitutive QS-cooperator is only weakly dependent on its frequency in the population (linear model: *α* = 0.389, *s. e.* = 0.066, *F*_1,6_ = 34.8, *P* < 0.001), whereas the positive-feedback QS-cooperator strongly reduces its cooperative investment as the frequency of response mutants increases (linear model: *β* = 1.159, *s. e.* = 0.086, *F*_1,6_ = 182, *P* < 0.0001). Results were not strongly affected by the choice of bacterial density where frequency-dependence was studied (Fig. S3). Gene-expression results therefore suggest that the positive-feedback QS-cooperator will be less exploited by response mutants and therefore better immune against them, as the theory predicts. The deviation of the constitutive QS-cooperator from constant (frequency-independent) response may depend on other QS systems involved in the activation of this system (see discussion).

### Swarming competitions verify the two predicted advantages of positive-feedbacks: reduced intra-group disadvantage and increased inter-group advantage

To further verify the theoretical predictions, we performed competitions of each cooperator against its corresponding response mutant in a swarming assay. Briefly, mixed cultures were plated in the middle of a semi-solid agar plate and allowed to swarm and cover the plate (see Methods for further details). Using a flow-cytometer we measured the relative fitness of cooperators after 2 days of growth on swarming plates. We sought an IPTG concentration, in which the two cooperators suffered a similar fitness decrease (cost) when co-cultured with a rare (1%) response mutant (i.e., when their cooperative investment is similar). This type of comparison was much more sensitive to small differences between the two cooperators than just comparing the swarming yield in mono-cultures (see discussion). IPTG dependence was different than that observed in liquid culture (Fig. 2D). This difference may be due to difference in bacterial densities and signal diffusion in the swarming assay. We found that in swarming conditions, the constitutive QS-cooperator suffered a stronger fitness cost compared to the positive-feedback QS-cooperator even with no IPTG. We therefore replaced the IPTG-inducible promoter with one that is less leaky (P_spac_ instead of P_hs_, see Methods). This allowed us to use inducer on the swarming plate and re-calibrate the IPTG level to compare the costs of constitutive and feedback QS-cooperators.

Fig. 3A presents the relative fitness of each of the two cooperators (now approximately comparable at 40μM IPTG) in competition with its respective response-null mutant in different initial frequency. The fitness cost of the constitutive QS-cooperator is only weakly dependent of its frequency in the population (linear model: *β* = −0.0015, *s. e.* = 0.0007, *F*_1,30_ = 4.28, *P* < 0.05), while the relative fitness of the positive-feedback QS-cooperator decreases strongly with the decrease in its frequency (linear model: *β* = −0.0109, *s. e.* = 0.0004, *F*_1,30_ = 895, *P* < 0.0001), as was suggested by the decrease in its cooperative investment (Fig. 2E). Interestingly, the positive-feedback QS-cooperator even wins as a minority, suggesting a preferential access to benefits that yields a snowdrift type of interaction [16] (see Discussion).

**Figure 3.**
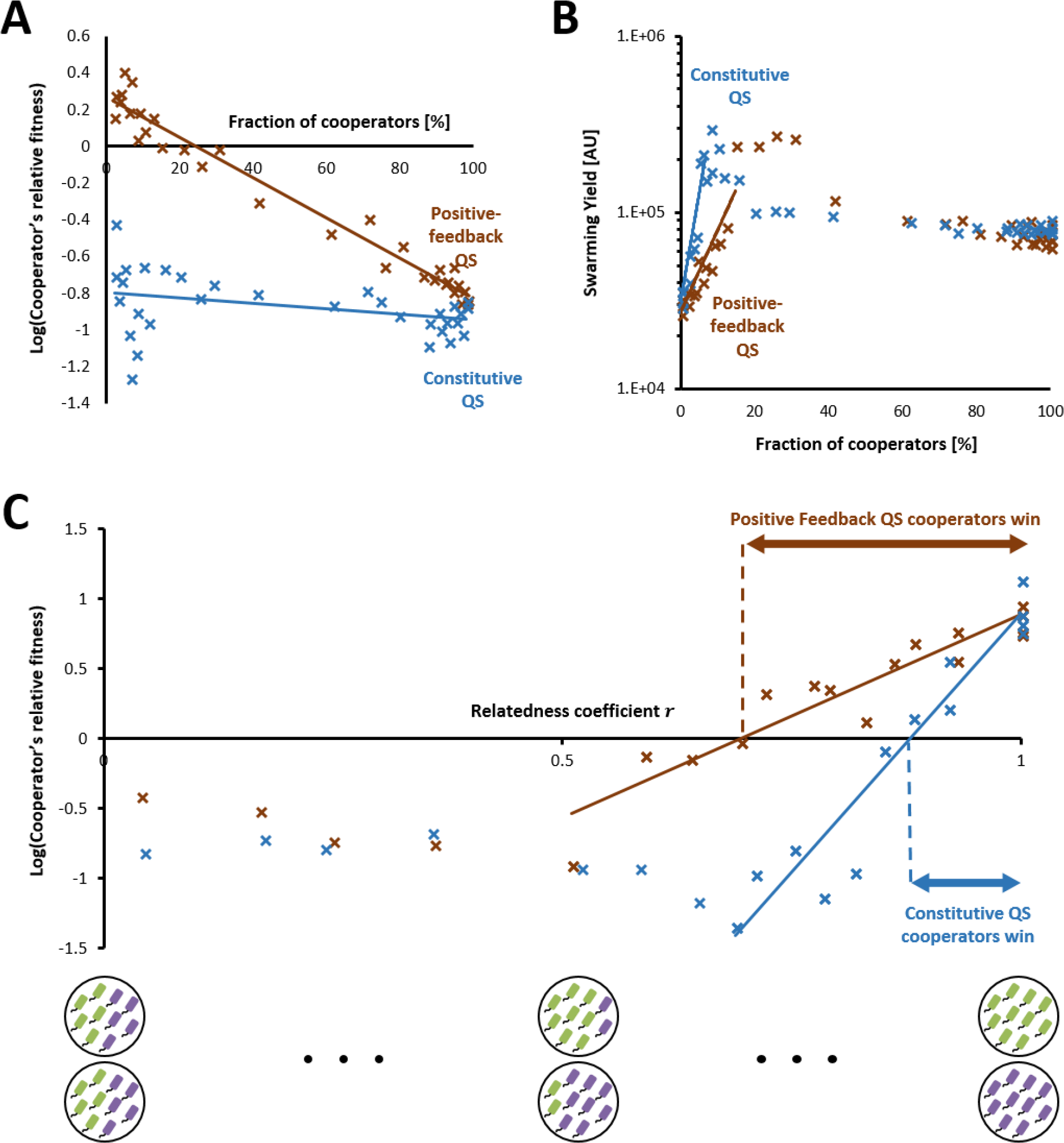
Swarming competitions between cooperators and their respective response-null mutants. Shown are **(A)** The logarithm of relative fitness of cooperators over response-null cheaters and **(B)** overall yield in swarming plates of co-cultures of constitutive QS-cooperators (AES3938, blue) or positive-feedback QS-cooperators (AES3634, brown) with their corresponding response-null mutants (AES3940 and AES3944, respectively). Each point describes a competition on one plate. Lines in (A) represent linear model fits to the data with *R*^2^ = 0.125 and *R*^2^ = 0.97 for constitutive and positive-feedback QS-cooperators respectively. Lines in (B) represent exponential fits to the increasing trend in the data with *R*^2^ = 0.89 and *R*^2^ = 0.83 for constitutive and positive-feedback QS-cooperators respectively. Results shown are for four biological repeats taken in different days for each co-cultured pair, which were used to start competitions at multiple initial frequencies. The IPTG concentrations for the constitutive QS-cooperator plates are 40μM. **(C)** Based on results shown in A,B, we calculated the relative fitness of the two types of cooperators in a structured population with different values of the relatedness coefficient *r*. The lines represent linear fits for part of the data where relative fitness increases, with *R*^2^ = 0.92, and *R*^2^ = 0.88 for constitutive QS-cooperators (blue), and positive-feedback QS-cooperators (brown), respectively. Each fit intersection with the X-axis is indicated and defines a threshold relatedness, above which cooperators win. The simple two-plate metapopulation analysis used to calculate the population structure is illustrated below the graph - combinations of two-plates with equal mean and increasing standard variation correspond to populations with increasing relatedness coefficient.

Fig. 3B shows that the overall growth on these plates sharply increases when increasing the initial frequency of both types of cooperators, with a gradual decrease at higher frequencies that is due to saturating benefit but increasing cost of surfactin production [17,60] (see Discussion for a possible explanation for this extreme saturation). While a very small frequency (~5%) of constitutive QS-cooperators is enough to yield substantial swarming, a significantly larger frequency (~15%) of positive-feedback QS-cooperators is needed for substantial swarming to take place. The intuitive reason for this is that when surrounded by their response mutants, the positive-feedback QS-cooperators produce less surfactin than constitutive QS-cooperators, as illustrated by the gene expression results (Fig. 2E). Fig. S4 presents similar comparisons between the two types of cooperators in swarming competitions, but with a lower concentration (27μM) of IPTG.

### Reanalysis in a simple structured-population model demonstrates the increased resistance of PFB against response mutants

The two comparisons – relative fitness and net yield – correspondingly represent the within- and between-group impact of cooperation on selection. Both type of interactions come into effect in structured population, as determined by the relatedness level *r*. To model this combined effect, we considered two-plate combinations from Fig. 3A,B (each plate is a well-mixed population, to a good approximation). We originally divided the two competing strains to pairs of swarming plates with opposite frequencies such that their combination yields a metapopulation with an overall frequency of 50-50. The relatedness coefficient *r* in group-structured populations is proportional to the variance in group composition (see Methods). In one extreme, a combination of two co-cultures with identical compositions yields a well-mixed population (*r* = 0), and in the other extreme, a combination of two mono-cultures (extreme variance in the composition of the two plates) yields a well-separated population (*r* = 1). Fig. 3C presents the relative fitness of the different cooperators in these two-plate meta-populations. For both types of cooperators there is a threshold relatedness, above which cooperators win, as expected from Hamilton’s rule. Furthermore, as predicted here theoretically, the threshold for the positive-feedback QS-cooperator is lower than that of the constitutive QS-cooperator, and thus the positive-feedback structure allows stable cooperation in a wider range of population structures. Again, using a different concentration of IPTG for comparing the two cooperators, yielded very similar results (Fig. S4).

## Discussion

In this work, we tested the effect of feedback loops in regulation of cooperative behaviors on their evolutionary stability. We theoretically showed that positive feedbacks on public goods extend the range of environments that maintain cooperation in the face of either null-mutant cheaters or small mutants of the regulatory pathway, and that this improvement in stability increases with feedback strength. This is due to a reduction in cooperative investment around cheaters that promotes the fitness of cooperators in group-structured populations in two ways. Firstly, positive-feedback cooperators suffer a smaller fitness cost in mixed groups with cheaters, compared to constitutive cooperators, thereby reducing cooperation’s intra-group disadvantage. Secondly, positive feedbacks reduce the average fitness of cheater-dominated groups, thereby improving the between-group advantage of cooperator-dominated groups.

Following the intuition of reduced cooperation around cheaters, we have also shown that these results generalize to other secreted molecules that control cooperative behaviors, e.g., QS signals. A simple QS design provides increased immunity to QS-null mutants, since cooperators can “sense” their presence and reduce cooperative investment. However, the simple QS design does not allow “sensing” of response-null mutants that have functional signaling genes and continue to produce QS signals and coerce QS-cooperators to cooperate strongly, even in cheater-dominated groups. Improved stability against these response-null mutants is only accomplished when there is a positive-feedback on the QS signal, as often occurs for QS systems [34,35].

This extension suggests a connection between public goods as regulators of cooperative behaviors to QS signals. If it is beneficial that public goods acquire some regulatory capability to activate their own transcription, this capability could also be used to regulate other cooperative behaviors. Indeed, some public goods have this capability, affording them a role of QS signals [27]. This could also suggest that some QS signals evolved from, and might be still active as public goods.

We have also tested the theoretical predictions experimentally. We used the ComQXPA QS system that controls, among other cooperative behaviors, the secretion of surfactin, a public-good molecule that enables swarming motility. We genetically modified its constitutive regulation, to either a positive-feedback design, or an IPTG-inducible constitutive design. This additional inducible system allowed us to meaningfully calibrate the open- and closed-loop designs to ensure similar cooperative investment of both cooperators in mono-cultures (as we verified with gene expression measurements, and fitness-cost measurements in cooperator-dominated co-cultures). This comparison is the only one that makes the two cooperators neutral when co-cultured together. Any difference in cooperative investment between them in mono-cultures, which must yield a difference in their co-culture, will only result in a fitness cost to one of them, while the benefit will be shared.

The gene-expression measurements verified that the cooperative investment of positive-feedback QS-cooperators decreased as their frequency in co-cultures with response-null mutants decreased. As a result, positive-feedback QS-cooperators should enjoy a two-fold advantage over constitutive QS-cooperators, i.e., a decreased fitness cost and a reduced yield in cheater-dominated co-cultures, as we demonstrated with swarming competitions. Furthermore, by re-analyzing competitions as two-plate meta-populations, we have also demonstrated how this advantage is translated into improved stability against response-null mutants in structured populations. As we predicted theoretically, positive-feedback QS-cooperators were stable against cheaters in a wider range of meta-populations, compared to constitutive QS-cooperators, thereby reducing the relatedness threshold required for stable cooperation.

Although the theoretical analyses were restricted to simple linear regulatory and fitness functions, the experimental support suggests that the essential components of the problem were indeed captured. Actually, the measured dependence of the cooperative investment of positive-feedback QS-cooperators on their frequency (Fig. 2E) is at the linear regime (*R*^2^ = 0.968, *n* = 8). More complex positive-feedback designs on public goods or QS signals could lead to bistability of the population as a whole (because of the extra-cellular nature of these molecules). This could reduce the cooperative investment of positive-feedback cooperators in cheater-dominated groups even further [61] but could also lead to hysteresis and more complex dynamics.

Another caveat of the theoretical analyses is the restriction to group-structured populations with constant group size, an assumption that was also made in the population structure analysis of experimental data. Taking population size into account, the positive-feedback structure would actually allow sensing of the density of cooperators (or signal secretors), and not their frequency in the population. This could affect the population dynamics and should be considered in future work. Apart from further theoretical investigation of the impact of group-size, it would be interesting to study the effect of feedbacks in more realistic population structures with simulations or experiments. However, in swarming, the increased population size due to migration, does not change population density significantly and therefore this is less relevant to the experimental analysis.

In a more specific manner, our swarming experiments revealed several surprising insights into *B. subtilis* swarming behavior. First, we found that the open-loop design also showed frequency-dependent activity and selection (Figs. 2E,3A) when mixed with response-null cheaters. This indicates the existence of a positive feedback loop in the native system. This may either result from a yet uncharacterized regulation of the ComQXP system, or from an effect of Rap-Phr quorum-sensing systems on ComA activity [18,62,63]. Second, we found that, at low frequency, positive-feedback cooperators had a fitness advantage over cheaters. This could either be explained by a semi-privatization of secreted surfactin molecules, leading to a snowdrift game interaction [16], or by population de-mixing [64,65] occurring in weak-swarming cheater-dominated plates. Interestingly, a similar phenomenon was observed for *P. aeruginosa* swarming [66]. Finally, we find that proficient swarming requires a very low frequency of the constitutive-QS strain (~5%) in a population of cheaters. This results from the very high level of surfactin produced by the swarming proficient derivatives of the lab strain, which has been selected for increased *srfA* expression and loss of surfactin production during domestication [17,60,67]. The wild strain produces less surfactin and would accordingly require high level of cooperation to allow for full swarming.

Altogether, our results point to a design principle of microbial cooperative behaviors. The improved stability against cheating, granted by these intercellular positive feedbacks, suggests one explanation for their abundance in regulatory structures of cooperative behaviors. Moreover, these results point to the positive-feedback regulatory structure as another microbial mechanism that may helped to maintain cooperative behaviors in evolution. These two complementary contributions of this project demonstrate the utility of combining social-evolutionary tools with the systems biology way of thinking.

## Methods

### Growth media and conditions

**Routine growth** was performed in Luria–Bertani [LB] broth (1% [w/v] tryptone (Difco), 0.5% [w/v] yeast extract (Difco), 0.5% [w/v] NaCl) or Spizizen minimal medium [SMM] (2 g L^−1^ (NH_4_)_2_SO_4_, 14 g L^−1^ K_2_HPO_4_, 6 g L^−1^ KH_2_PO_4_, 1 g L^−1^ disodium citrate, 0.2 g L^−1^ MgSO_4_·7H_2_O). SMM media was supplemented with 0.5% [w/v] glucose and trace elements (125 mg L^−1^ MgCl_2_·6H_2_O, 5.5 mg L^−1^ CaCl_2_, 13.5 mg L^−1^ FeCl_2_·6H_2_O, 1 mg L^−1^ MnCl_2_·4H_2_O, 1.7 mg L^−1^ ZnCl_2_, 0.43 mg L^−1^ CuCl_2_·4H_2_O, 0.6 mg L^−1^ CoCl_2_·6H_2_O, 0.6 mg L^−1^ Na_2_MoO_4_·2H_2_O). Isopropyl β-D-thiogalactopyranoside (IPTG, Sigma) was added to the medium at the indicated concentration when appropriate. 0.01 M Phosphate buffer saline (PBS) pH 7.4 (Sigma) was used for dilution and suspension of cells. Petri dishes for routine procedures were solidified using 2% [w/v] agar (Difco). Antibiotic concentrations: Macrolides-Lincosamides-Streptogramin B (MLS, 1 μg ml^−1^ Erythromycin, 25 μg ml^−1^ Lincomycin); Spectinomycin (Sp, 100 μg ml^−1^); Tetracycline (Tet, 10 μg ml^−1^); Kanamycin (Kan, 10 μg ml^−1^); Chloramphenicol (Cm, 5 μg ml^−1^); Ampicillin (Amp, 100 μg ml^−1^), Phleomycin (Ph, 2.5 μg ml^−1^).

#### Pre-measurement growth protocol

Prior to all measurements, an overnight colony from an LB agar plate was inoculated in 1.5 mL SMM liquid medium and grown (with shaking) for ~7 hours in 37 °C until an OD_600_ of 0.1-0.3 was reached. The cultures were then diluted by a factor of 10^6^ and grown (with shaking) overnight at 37°C. We find that this long incubation in minimal medium both reduced the effects of QS prior to growth and reduced the arbitrary difference in growth between two co-cultured wild-type colonies.

### Gene expression experiments

In all experiments, the overnight cultures from the pre-measurement growth protocol, were measured in the following day. Samples were taken at several time points and the OD_600_-equivalent (see below) and YFP levels were measured by flow cytometry (see below). YFP level was determined from the median level of the unimodal distribution of YFP expressing cells using flow cytometry (see below). YFP level was normalized by the auto-fluorescence of the wild-type. The expected YFP levels at the chosen bacterial density were calculated by either interpolation of two consecutive measurements, one below and one above the chosen OD_600_, if existed, or by extrapolation of two consecutive measurements from the same side of the chosen bacterial density.

#### OD – flow-cytometer calibration

We used the wild type strain to calibrate flow-cytometer density measurement to OD_600_ measurement. At several time points during growth, the number of flow-cytometer events in 60 seconds (considering the dilution factor in the flow-cytometer sample preparation), the flow-cytometer flow rate (using Flow-check Pro Fluorophores, Beckman-Coulter), and OD_600_ were measured. A calibration factor of 220,000 events per OD_600_ = 0.01 was obtained from the slope of the linear fit to the calibration curve. At all subsequent experiments, the flow-cytometer flow rate was re-measured in the same way to correct for day to day variation in the flow rate.

#### For co-culture experiments

Cells of different strains were mixed in appropriate ratios, based on relative optical density, before the overnight part of the pre-measurement growth protocol. The exact ratios were measured, together with overall density and YFP, in the following day using flow cytometry. Both strains carried the QS-response YFP transcriptional reporter, and the YFP-measured strain had an additional mCherry constitutive marker.

#### Flow cytometry

Samples were analyzed on a Gallios flow-cytometer (Beckman-Coulter), equipped with 4 lasers (405 nm, 488 nm co-linear with 561 nm, and 638 nm). The emission filters used were: YFP – 525/40, mCherry – 620/30. Events were discriminated using the forward-scatter parameter. For each run, discrimination enabled a single, well-defined population to appear in the forward-scatter (FS) by side-scatter plot. Gating on the fluorescent populations and inspection of the non-discriminated forward by side-scatter plot indicated that over 99.9% of the fluorescent cells are present in the discriminated population. In all analyzed samples, only single cells were considered by removing FS events, whose time-of-flight was correlated to the integral of the signal. Gating of the different fluorescent populations was performed by inspection of the log-log FLx by FLy plots (x & y represent the appropriate filter for each fluorescent marker), where two distinct populations were clearly visible, resulting in a type-I and type-II errors of less than 0.05%. For each run, at least 100,000 cells were analyzed, and the total events analyzed such that the minority population was never below 1,000 events.

### Swarming competition experiments

Cells were grown as described in the pre-measurement growth protocol, and were then centrifuged, re-suspended in PBS, and diluted to an OD_600_ of 0.01. Agar plates (0.8%) containing 25 mL of SMM medium supplemented with trace elements, 0.03% glucose, and IPTG (at the indicated concentration when appropriate) were poured, at a constant temperature, in a laminar flow chamber and left there to solidify for 24 minutes. Five microliters of the diluted cultures were consequently placed at the centers of the plates, which were then left for 11 minutes in the laminar flow chamber, for the five-microliter drop to dry. The plates were then incubated at 30°C for 47 hours, after which the swarms were scraped and collected from the plates by suspension in 2 ml of PBS. The final densities (OD_600_-equivalent) of both strain were measured by flow cytometry as described above, from the two distinct sub-populations, and compared to the initial densities, measured at time zero. The fitness (growth factor) of each strain was calculated as the ratio of final and initial densities, multiplied by the ratio of final and initial liquid volumes (5μl and 2ml respectively). The cooperator’s relative fitness is the ratio between the cooperator’s and the cheater’s fitness, while the overall swarming yield is the weighted average of fitness of the two strains. In all competitions, both cooperators and cheaters carried a constitutive YFP reporter, while cheaters carried an additional constitutive mCherry reporter (see strain list in Table S1). It was shown previously [17] that the additional mCherry reporter does not carry a significant fitness cost, as we also verified it with these strains.

### Analysis in structured populations

To simulate population structure, we originally divided the two competing strains to pairs of swarming plates with opposite frequencies (i.e., if in one plate the frequency *G* of cooperators was *G* = *x*, then in the second plate their frequency *G* was approximately *G* = 1 − *x*, such that the frequency of cooperators in the meta population composed of the two plates was approximately 〈*G*〉~0.5). The relatedness of the two-plate populations was calculated using the following formula [49]:

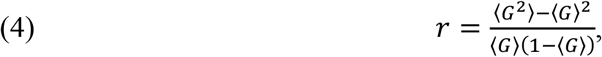

where 〈 〉 denotes an average over the two plates, each weighted by its overall density (OD_600_-equivalent). The fitness (growth factor) of each strain in the two-plate population was calculated as described above for single plates but using its densities (initial and final) in the overall two-plate population. The relative fitness of cooperators is again the ratio between the cooperator’s fitness and the cheater’s fitness.

### Strain construction

All *B. subtilis* and *E. coli* strains are detailed in Table S1 and Table S2 respectively, while respective primers are provided in Table S3. Deletion mutations and their replacement with the indicated antibiotic resistance cassette were performed using the long flanking homology PCR methods [68]. 1kb fragments corresponding to regions upstream and downstream of the target gene were amplified by PCR. The 5’ end of the reverse primer for the upstream region and the 3’ end of the forward primer for the downstream region contained a short overhang sequence homologous to an antibiotic resistance cassette. A second PCR was then performed using the PCR products of the first reaction as primers, and the antibiotic cassette as a template. The final product was transformed to *B. subtilis* PY79 (AES101). Integration of the construct to the genomic DNA was confirmed by PCR. Accordingly, *comXP* and *comXP-comA* were deleted from the PY79 chromosome using the primers comXP-del-P1-P4 and comXPA-del-P1-P4 (Table S3).

The mutations and constructs were transferred to PY79 either by natural transformation [69] or by SPP1-mediated generalized transduction [70]. Integration of *amyE* integration plasmids into the *zjd89::amyEΩ Cm* Kan [71] was done in two steps. First, the plasmid was integrated into a PY79 strain (AES2864) carrying the zjd89 construct and was screened for an Amy+ phenotype. A SPP1 lysate [70] of the resulting strain was then inserted into other strains with selection for either Kan or Cm, depending on the genetic background of the integrated genome.

Construction of *zjd-89*::(P_hs_-*comXP* Kan Cm Sp) was performed by PCR amplification of *comXP*, using AES101 as a template and the comXP-F/comXP-R primer pair (Table S3). The PCR fragment was digested with NheI and SphI and ligated downstream of the hyperspank promoter (P_hs_) of the pDR111 vector containing Sp resistance.

Construction of *zjd-89*::(P_srfA_-*comXP* Kan Cm Sp) was performed by restricting of the *srfA* promoter (P_srfA_) from AEC945 with EcoRI and NheI, and subcloning it into pDR111::P_hs_-comXP plasmid (AEC1054) instead of the hyperspank promoter (P_hs_).

Construction of *zjd-89*::(P_spac_-*comXP* Kan Cm Sp) was performed by PCR amplification of the spac promoter (P_spac_), using BD1916 as a template and the Pspac-F/ Pspac-R primer pair (Table S3). The PCR fragment was digested with EcoRI and NheI and ligated into pDR111::P_hs_-comXP plasmid (AEC1054) instead of the hyperspank promoter (P_hs_).

## References

1. Alon U (2007) Network motifs: Theory and experimental approaches. Nat Rev Genet 8: 450–461.

2. Liebermeister W, Klipp E, Schuster S, Heinrich R (2004) A theory of optimal differential gene expression. BioSystems 76: 261–278.

3. Ben-Zvi D, Barkai N (2010) Scaling of morphogen gradients by an expansion-repression integral feedback control. Proc Natl Acad Sci 107: 6924–6929.

4. Yi T-M, Huang Y, Simon MI, Doyle J (2000) Robust perfect adaptation in bacterial chemotaxis through integral feedback control. Proc Natl Acad Sci 97: 4649–4653.

5. Dublanche Y, Michalodimitrakis K, Kümmerer N, Foglierini M, Serrano L (2006) Noise in transcription negative feedback loops: Simulation and experimental analysis. Mol Syst Biol 2: 41.

6. Rosenfeld N, Elowitz MB, Alon U (2002) Negative autoregulation speeds the response times of transcription networks. J Mol Biol 323: 785–793.

7. Ozbudak EM, Thattal M, Lim HH, Shraiman BI, Van Oudenaarden A (2004) Multistability in the lactose utilization network of Escherichia coli. Nature 427: 737–740.

8. Venturelli OS, El-Samad H, Murray RM (2012) Synergistic dual positive feedback loops established by molecular sequestration generate robust bimodal response. Proc Natl Acad Sci U S A 109: E3324–33.

9. Isaacs FJ, Hasty J, Cantor CR, Collins JJ (2003) Prediction and measurement of an autoregulatory genetic module. Proc Natl Acad Sci 100: 7714–7719.

10. Ferrell JE (2002) Self-perpetuating states in signal transduction: Positive feedback, double-negative feedback and bistability. Curr Opin Cell Biol 14: 140–148.

11. Mitrophanov AY, Groisman EA (2008) Positive feedback in cellular control systems. BioEssays 30: 542–555.

12. West SA, Griffin AS, Gardner A, Diggle SP (2006) Social evolution theory for microorganisms. Nat Rev Microbiol 4: 597–607.

13. Celiker H, Gore J (2013) Cellular cooperation: Insights from microbes. Trends Cell Biol 23: 9–15.

14. Diggle SP, Griffin AS, Campbell GS, West SA (2007) Cooperation and conflict in quorum-sensing bacterial populations. Nature 450: 411–414.

15. Sandoz KM, Mitzimberg SM, Schuster M (2007) Social cheating in Pseudomonas aeruginosa quorum sensing. Proc Natl Acad Sci 104: 15876–15881.

16. Gore J, Youk H, Van Oudenaarden A (2009) Snowdrift game dynamics and facultative cheating in yeast. Nature 459: 253–256.

17. Pollak S, Omer-Bendori S, Even-Tov E, Lipsman V, Bareia T, Ben-Zion I, Eldar A (2016) Facultative cheating supports the coexistence of diverse quorum-sensing alleles. Proc Natl Acad Sci 113: 2152–2157.

18. Even-Tov E, Omer Bendori S, Valastyan J, Ke X, Pollak S, Bareia T, Ben-Zion I, Bassler BL, Eldar A (2016) Social Evolution Selects for Redundancy in Bacterial Quorum Sensing. PLoS Biol 14: e1002386.

19. Xavier JB, Kim W, Foster KR (2011) A molecular mechanism that stabilizes cooperative secretions in Pseudomonas aeruginosa. Mol Microbiol 79: 166–179.

20. Venturi V, Bertani I, Kerényi Á, Netotea S, Pongor S (2010) Co-swarming and local collapse: Quorum sensing conveys resilience to bacterial communities by localizing cheater mutants in Pseudomonas aeruginosa. PLoS One 5: e9998.

21. West SA, Buckling A (2003) Cooperation, virulence and siderophore production in bacterial parasites. Proc R Soc B Biol Sci 270: 37–44.

22. Diard M, Garcia V, Maier L, Remus-Emsermann MNP, Regoes RR, Ackermann M, Hardt WD (2013) Stabilization of cooperative virulence by the expression of an avirulent phenotype. Nature 494: 353–356.

23. Raymond B, West SA, Griffin AS, Bonsall MB (2012) The dynamics of cooperative bacterial virulence in the field. Science (80-) 336: 85–88.

24. Schuster M, Greenberg EP (2006) A network of networks: Quorum-sensing gene regulation in Pseudomonas aeruginosa. Int J Med Microbiol 296: 73–81.

25. Takezawa M, Price ME (2010) Revisiting ‘The Evolution of Reciprocity in Sizable Groups’: Continuous reciprocity in the repeated n-person prisoner’s dilemma. J Theor Biol 264: 188–196.

26. Poole K (2003) Iron acquisition and its control in pseudomonas aeruginosa many roads lead to rome. Front Biosci 8: 1051.

27. Lamont IL, Beare PA, Ochsner U, Vasil AI, Vasil ML (2002) Siderophore-mediated signaling regulates virulence factor production in Pseudomonas aeruginosa. Proc Natl Acad Sci 99: 7072–7077.

28. Brickman TJ, Ho Young Kang, Armstrong SK (2001) Transcriptional activation of Bordetella alcaligin siderophore genes requires the AlcR regulator with alcaligin as inducer. J Bacteriol 183: 483–489.

29. Ambrosi C, Leoni L, Visca P (2002) Different responses of pyoverdine genes to autoinduction in Pseudomonas aeruginosa and the group Pseudomonas fluorescens-Pseudomonas putida. Appl Environ Microbiol 68: 4122–4126.

30. Reimmann C, Serino L, Beyeler M, Haas D (1998) Dihydroaeruginoic acid synthetase and pyochelin synthetase, products of the pchEF genes, are induced by extracellular pyochelin in Pseudomonas aeruginosa. Microbiology 144: 3135–3148.

31. Pugsley AP (1983) Autoinduced synthesis of colicin E2. MGG Mol Gen Genet 190: 379–383.

32. Ghazaryan L, Soares MIM, Gillor O (2014) Auto-regulation of DNA degrading bacteriocins: Molecular and ecological aspects. Antonie van Leeuwenhoek, Int J Gen Mol Microbiol 105: 823–834.

33. Mavridou DAI, Gonzalez D, Kim W, West SA, Foster KR (2018) Bacteria Use Collective Behavior to Generate Diverse Combat Strategies. Curr Biol 28: 345–355.e4.

34. Ng W-L, Bassler BL (2009) Bacterial Quorum-Sensing Network Architectures. Annu Rev Genet 43: 197–222.

35. Allen RC, McNally L, Popat R, Brown SP (2016) Quorum sensing protects bacterial co-operation from exploitation by cheats. ISME J 10: 1706–1716.

36. Tanouchi Y, Tu D, Kim J, You L (2008) Noise reduction by diffusional dissipation in a minimal quorum sensing motif. PLoS Comput Biol 4: e1000167.

37. Tabareau N, Slotine JJ, Pham QC (2010) How synchronization protects from noise. PLoS Comput Biol 6: e1000637.

38. Weber M, Buceta J (2011) Noise regulation by quorum sensing in low mRNA copy number systems. BMC Syst Biol 5: 11.

39. Ross-Gillespie A, Kümmerli R (2014) Collective decision-making in microbes. Front Microbiol 5:.

40. Williams JW, Cui X, Levchenko A, Stevens AM (2008) Robust and sensitive control of a quorum-sensing circuit by two interlocked feedback loops. Mol Syst Biol 4: 234.

41. Haseltine EL, Arnold FH (2008) Implications of rewiring bacterial quorum sensing. Appl Environ Microbiol 74: 437–445.

42. Stabb E V. (2018) Could positive feedback enable bacterial pheromone signaling to coordinate behaviors in response to heterogeneous environmental cues? MBio 9: e00098–18.

43. Gilbert OM, Foster KR, Mehdiabadi NJ, Strassmann JE, Queller DC (2007) High relatedness maintains multicellular cooperation in a social amoeba by controlling cheater mutants. Proc Natl Acad Sci 104: 8913–8917.

44. Dandekar AA, Chugani S, Greenberg EP (2012) Bacterial quorum sensing and metabolic incentives to cooperate. Science (80-) 338: 264–266.

45. Wang M, Schaefer AL, Dandekar AA, Greenberg EP (2015) Quorum sensing and policing of *Pseudomonas aeruginosa* social cheaters. Proc Natl Acad Sci 112: 2187–2191.

46. Schluter J, Schoech AP, Foster KR, Mitri S (2016) The Evolution of Quorum Sensing as a Mechanism to Infer Kinship. PLoS Comput Biol 12: e1004848.

47. Brown SP (1999) Cooperation and conflict in host-manipulating parasites. Proc R Soc B Biol Sci 266: 1899–1904.

48. Brown SP, Johnstone RA (2001) Cooperation in the dark: signalling and collective action in quorum-sensing bacteria. Proc R Soc B Biol Sci 268: 961–965.

49. Pepper JW (2000) Relatedness in trait group models of social evolution. J Theor Biol 206: 355–368.

50. Gardner A, West SA, Wild G (2011) The genetical theory of kin selection. J Evol Biol 24: 1020–1043.

51. Kearns DB, Losick R (2003) Swarming motility in undomesticated Bacillus subtilis. Mol Microbiol 49: 581–590.

52. Lopez D, Fischbach MA, Chu F, Losick R, Kolter R (2009) Structurally diverse natural products that cause potassium leakage trigger multicellularity in Bacillus subtilis. Proc Natl Acad Sci 106: 280–285.

53. Das P, Mukherjee S, Sen R (2008) Genetic regulations of the biosynthesis of microbial surfactants: An overview. Biotechnol Genet Eng Rev 25: 165–186.

54. James BL, Kret J, Patrick JE, Kearns DB, Fall R (2009) Growing bacillus subtilis tendrils sense and avoid each other: Research letter. FEMS Microbiol Lett 298: 12–19.

55. Kearns DB (2010) A field guide to bacterial swarming motility. Nat Rev Microbiol 8: 634–644.

56. Nakano MM, Magnuson R, Myers A, Curry J, Grossman AD, Zuber P (1991) srfA is an operon required for surfactin production, competence development, and efficient sporulation in Bacillus subtilis. J Bacteriol 173: 1770–1778.

57. Grossman AD (1995) Genetic Networks Controlling the Initiation of Sporulation and the Development of Genetic Competence in Bacillus subtilis. Annu Rev Genet 29: 477–508.

58. Weinrauch Y, Penchev R, Dubnau E, Smith E, Dubnau D (1990) A Bacillus subtilis regulatory gene product for genetic competence and sporulation resembles sensor protein members of the bacterial two-component signal-transduction systems. Genes Dev 4: 860–872.

59. Ohsawa T, Tsukahara K, Sato T, Ogura M (2006) Superoxide stress decreases expression of srfA through inhibition of transcription of the comQXP quorum-sensing locus in Bacillus subtilis. J Biochem 139: 203–211.

60. Omer Bendori S, Pollak S, Hizi D, Eldar A (2015) The RapP-PhrP quorum-sensing system of Bacillus subtilis strain NCIB3610 affects biofilm formation through multiple targets, due to an atypical signal-insensitive allele of RapP. J Bacteriol 197: 592–602.

61. Cavaliere M, Poyatos JF (2013) Plasticity facilitates sustainable growth in the commons. J R Soc Interface 10: 20121006.

62. Even-Tov E, Omer Bendori S, Pollak S, Eldar A (2016) Transient Duplication-Dependent Divergence and Horizontal Transfer Underlie the Evolutionary Dynamics of Bacterial Cell–Cell Signaling. PLOS Biol 14: e2000330.

63. Auchtung JM, Lee CA, Grossman AD (2006) Modulation of the ComA-dependent quorum response in Bacillus subtilis by multiple Rap proteins and Phr peptides. J Bacteriol 188: 5273–5285.

64. Hallatschek O, Hersen P, Ramanathan S, Nelson DR (2007) Genetic drift at expanding frontiers promotes gene segregation. Proc Natl Acad Sci 104: 19926–19930.

65. Van Dyken JD, Müller MJI, MacK KML, Desai MM (2013) Spatial population expansion promotes the evolution of cooperation in an experimental prisoner’s dilemma. Curr Biol 23: 919–923.

66. De Vargas Roditi L, Boyle KE, Xavier JB (2013) Multilevel selection analysis of a microbial social trait. Mol Syst Biol 9: 684.

67. Pollak S, Omer Bendori S, Eldar A (2015) A complex path for domestication of B. subtilis sociality. Curr Genet 61: 493–496.

68. Wach A (1996) PCR-synthesis of marker cassettes with long flanking homology regions for gene disruptions in S. cerevisiae. Yeast 12: 259–265.

69. Harwood CR, Cutting SM (1990) Molecular biological methods for Bacillus (Modern Microbiological Methods). Wiley.

70. Yasbin RE, Young FE (1974) Transduction in Bacillus subtilis by bacteriophage SPP1. J Virol 14: 1343–1348.

71. Vin YY, Kondratiev DE (2012) Information approach to studying byzantine law and its receptions. Byzantinoslavica 70: 76–96.

