## Supplementary Information for "Intercellular positive-feedback loops promote the evolutionary stability of microbial cooperative behaviors"

### Contents

**SUPPLEMENTARY TEXT** 2

POSITIVE FEEDBACKS ON PUBLIC GOODS INCREASE THE RANGE OF POPULATION STRUCTURES IN WHICH COOPERATION IS EVOLUTIONARILY STABLE TO SMALL MUTATIONS 2

GLOBAL INVASION ANALYSIS SHOWS THAT A POSITIVE-FEEDBACK STRUCTURE IMPROVES THE STABILITY AGAINST PUBLIC-GOOD NULL MUTANTS 5

APPLICATION TO ANOTHER EXAMPLE OF INTERCELLULAR FEEDBACKS – QUORUM SENSING REGULATION 6

**SUPPLEMENTARY TABLES** 8

**SUPPLEMENTARY REFERENCES** 11

**SUPPLEMENTARY FIGURES** 12

### Supplementary text

#### Positive feedbacks on public goods increase the range of population structures in which cooperation is evolutionarily stable to small mutations

A common approach to finding optimal strategies is adaptive dynamics theory, which deals with continuous traits that evolve through small phenotypic changes [1]. Using adaptive dynamics, we tackled the problem of evolutionary optimality of regulatory networks in a social context. We considered regulatory networks that are determined by a set of genes with varying values (a set of gene values is considered a genotype, forming a strategy), and searched among them for a locally evolutionarily-stable strategy (local ESS, a strategy that if adopted by all members of the population, is uninvasible by close strategies [2]).

To this aim, we used the simplest model of feedback-regulated cooperation, where a gene for a public-good molecule is linearly regulated by the average concentration of public goods in the surrounding environment, such that an individual's public-good production rate is:

(S1) $$x = f(\vec{\tilde{g}}, PG) = \tilde{g}_0 + \tilde{g}_{FB} PG,$$

where $x$ denotes the individual bacterium public-good production rate, $PG$ is the public-good concentration in the medium, and $\vec{\tilde{g}} = (\tilde{g}_0, \tilde{g}_{FB})$ is a set of values for the genes that determine the form of the regulatory function $f$. Specifically, $\tilde{g}_0 > 0$ is the basal public-good production rate, and $\tilde{g}_{FB}$ is the feedback strength that measures the effect of the public good on its own production rate).

The ordinary differential equation (ODE) for the number of public-good molecules, $PG$, in some constant volume occupied by a well-mixed bacterial population, is:

(S2) $$\frac{dPG}{dt} = \sum_{i=1}^{N} x_i - \gamma PG,$$

where $x_i = \tilde{g}_{0,i} + \tilde{g}_{FB,i} PG$ is the public-good production rate of bacterium $i$, $N$ is the assumingly constant number of bacteria, $\vec{\tilde{g}}_i = (\tilde{g}_{0,i}, \tilde{g}_{FB,i})$ is the strategy of individual $i$, and $\gamma$ is the public-good degradation rate. In the following, we assumed that protein expression dynamics is much faster than the evolutionary dynamics, and thus we could ignore the equilibration period and consider the steady state of public-good concentration, which is proportional to the population average production rate:

(S3) $$PG_{st} = \frac{N}{\gamma}\bar{x} = \frac{1}{\gamma}\sum_{i=1}^{N} x_i.$$

The individual strategy can therefore be written as:

(S4) $$x = f(\vec{g}, \bar{x}) = g_0 + g_{FB}\bar{x},$$

where $g_0 = \tilde{g}_0$ and $g_{FB} = \tilde{g}_{FB}\frac{N}{\gamma}$.

If the population is monomorphic ($\vec{g}_i = \vec{g}$ for all $i$), Eq. (S2) becomes:

(S5) $$\frac{dPG}{dt} = N(g_0 + g_{FB}\bar{x}) - N\bar{x},$$

and the steady state is thus:

(S6) $$\bar{x} = \frac{\gamma}{N} PG_{st} = \frac{g_0}{1-g_{FB}}.$$
We therefore restricted the analysis to $g_{FB} < 1$.

If the population is dimorphic, with two genotypes at frequencies $p, 1-p$, Eq. (S2) becomes:

(S7) $$\frac{dPG}{dt} = N\left(p\left(g_{0,1} + g_{FB,1}\bar{x}\right) + (1-p)\left(g_{0,2} + g_{FB,2}\bar{x}\right)\right) - N\bar{x},$$
and the steady-state mean public-good production rate follows:

(S8) $$\bar{x} = \frac{\gamma}{N} PG_{st} = \frac{p g_{0,1} + (1-p) g_{0,2}}{1 - p g_{FB,1} - (1-p) g_{FB,2}}.$$

To find ESSs, we considered the invasion of infinitesimally rare mutants into a population dominated by a resident genotype. The relatedness of an invader to its interaction group, in such a scheme, equals to the mean probability of invaders to find invaders in their group [3]. We therefore assumed a simple population structure such that the frequency of invaders (and not only its mean) in any group that is not purely of the resident genotype equals the relatedness coefficient $r$ [4,5].

We then looked for local ESSs (in the adaptive-dynamics, and not the original, definition) – strategies $\vec{g}^*$ that if adopted by all the population, are uninvasible by close strategies [2]. Thus, in such a population structure, a local ESS $\vec{g}^*$ must satisfy the following condition for every strategy $\vec{g}$ that is close to $\vec{g}^*$; The fitness of an individual of the invading strategy $\vec{g}$ in a mixed patch (with a frequency $r$ of invaders, the rest carry the ESS $\vec{g}^*$) should be lower than the fitness of a resident in a resident patch [2,4,5]:

(S9) $$W\left(x(\vec{g}; \vec{g}^*), \bar{x}(\vec{g}; \vec{g}^*)\right) < W\left(x(\vec{g}^*; \vec{g}^*), \bar{x}(\vec{g}^*; \vec{g}^*)\right),$$
where the first argument in $x$ and $\bar{x}$ is the individual strategy and the second argument is the resident strategy. Therefore $\vec{g} = \vec{g}^*$ is a maximum point of $W\left(x(\vec{g}; \vec{g}^*), \bar{x}(\vec{g}; \vec{g}^*)\right)$ and thus $\vec{g}^*$ could be found from the first order equations [2,4,5]:

(S10) $$\left.\frac{\partial W\left(x(\vec{g};\vec{g}^*), \bar{x}(\vec{g};\vec{g}^*)\right)}{\partial \vec{g}}\right|_{\vec{g}=\vec{g}^*} = \vec{0}.$$
Substituting $r$ instead of $p$ in Eq. (S8), yields the mean public-good production rate in such a patch:

(S11) $$\bar{x}(\vec{g}; \vec{g}^*) = \frac{r g_0 + (1-r) g_0^*}{1 - r g_{FB} - (1-r) g_{FB}^*},$$

while the invader's level of cooperation in this patch is:

(S12) $$x(\vec{g}; \vec{g}^*) = g_0 + g_{FB}\bar{x} = \frac{g_0 + (1-r)(g_0^* g_{FB} - g_0 g_{FB}^*)}{1 - r g_{FB} - (1-r) g_{FB}^*}.$$
We assumed that the fitness function $W$ depends only indirectly on $\vec{g}, \vec{g}^*$ through $x, \bar{x}$, and thus the first order equations can be written as:

(S13) $$\begin{cases} 0 = \left.\frac{\partial W(x(\vec{g};\vec{g}^*),\bar{x}(\vec{g};\vec{g}^*))}{\partial g_0}\right|_{\vec{g}=\vec{g}^*} = \left[\frac{\partial W}{\partial x}\frac{\partial x}{\partial g_0} + \frac{\partial W}{\partial \bar{x}}\frac{\partial \bar{x}}{\partial g_0}\right]_{\vec{g}=\vec{g}^*} \\ 0 = \left.\frac{\partial W(x(\vec{g};\vec{g}^*),\bar{x}(\vec{g};\vec{g}^*))}{\partial g_{FB}}\right|_{\vec{g}=\vec{g}^*} = \left[\frac{\partial W}{\partial x}\frac{\partial x}{\partial g_{FB}} + \frac{\partial W}{\partial \bar{x}}\frac{\partial \bar{x}}{\partial g_{FB}}\right]_{\vec{g}=\vec{g}^*} \end{cases}.$$

We first assumed a linear fitness function, where the fitness of each individual decreases with its own public-good production rate (with a cost parameter $C$) and increases with the mean public-good production rate in its surroundings (with a benefit parameter $B$)

(S14) $$W(x,\bar{x}) = 1 - Cx + B\bar{x}.$$

We assumed $B > C$, to have a monomorphic cooperator $(x > 0)$ population do better than a monomorphic cheater $(x = 0)$ population, as usually assumed in models of cooperation [6]. In a monomorphic population with no conflict ($x = \bar{x}$), the optimal public-good production rate is infinite. In the case of ESS analysis, Eqs. (S13) yield only one independent equation that sets a line of potential ESSs: $\vec{g}^* = \left(g_0^*\,,\ g_{FB}^* = \frac{1-\frac{B}{C}r}{1-r}\right)$, for which the second-derivative test is inconclusive. A test of nearby points demonstrated that these are not local maxima, but that the whole surface is flat, which results from the linearity of the fitness function.

We therefore considered a multiplicative-linear fitness form [4,5]:

(S15) $$W(x,\bar{x}) = (1 - Cx)(1 + B\bar{x}),$$

where we restricted the cooperation level (public-good production rate) to the region $x < \frac{1}{C} - \frac{1}{B}$ to have cooperators alone do better than cheaters alone. In a monomorphic population, the optimal level of public-good production is therefore:

(S16) $$x_{max} = \frac{1}{2C} - \frac{1}{2B},$$

so cooperation level could be further restricted to the monotonic region below this maximum.

Again, Eqs. (S13) yield only one independent equation, which results in the following relation between basal public-good production, feedback strength and relatedness:

(S17) $$g_0^* = \frac{1-g_{FB}^*}{B} \cdot \frac{r\left(\frac{B}{C}-g_{FB}^*\right)-(1-g_{FB}^*)}{1+r-g_{FB}^*(1-r)}.$$

For every value of the relatedness $0 < r < 1$, Eq. (S17) describes a curve of potential ESSs in the plane. That is, for every $g_{FB}^* < 1$ there is a corresponding $g_0^*$ and a corresponding (potential) ESS level of cooperation:

(S18) $$x^* = x(\vec{g}^*;\vec{g}^*) = \frac{g_0^*}{1-g_{FB}^*}.$$

A simple analysis shows that for every $r$, there is a range of $g_{FB}^*$: $g_{FB}^* < \frac{1-r\frac{B}{C}}{1-r}$, for which Eq. (S17) yields a negative $g_0^*$ and therefore negative $x^*$ (which is not allowed). It is easy to show numerically that for those values of $g_{FB}^*$, the strategy $(g_0 = 0, g_{FB} =$

$g_{FB}^*$), which represents zero cooperation level ($x^* = 0$, i.e., a cheater's strategy) is evolutionarily stable (we used Eq. (S9) to show that strategies in the vicinity of these strategies cannot invade it). It is worth noting that the strategy ($g_0 = 0, g_{FB} = g_{FB}^*$) is a cheater in monoculture (there is no basal public-good production, so feedback is irrelevant), but can be coerced to cooperate in co-culture with another cooperative strategy (with $g_0 > 0$).

On the other hand, we have shown numerically (using Eq. (S9)) that the potential ESSs ($g_0^*, g_{FB}^*$), obtained from Eq. (S17), that yield a positive cooperation level (i.e., for $g_{FB}^* > \frac{1-r\frac{B}{C}}{1-r}$) are indeed local maxima. Therefore, for each relatedness $r$ the ESS curve is made of two parts (Fig. S1), one which corresponds to an evolutionarily-stable cooperative strategy (i.e., $g_0^* > 0$ obtained from Eq. (S17)), and one at which $g_0^* = 0$ (i.e., cheating is evolutionarily stable).

Another way to describe this result is that an ESS ($g_0^*, g_{FB}^*$) represents a positive level of cooperation, only when the relatedness is higher than a threshold that depends on $g_{FB}^*$:

(S19) $$r > r_{th}(g_{FB}^*) = \frac{1-g_{FB}^*}{\frac{B}{C}-g_{FB}^*}.$$

Fig. S2 shows this threshold (red line) as a function of the feedback strength $g_{FB}^*$. The blue dot is the Hamilton rule threshold for constitutive cooperation: $r_{th}(g_{FB}^* = 0) = \frac{C}{B}$. One can see that as $g_{FB}^*$ becomes larger, the range of relatedness levels that allow a positive evolutionarily-stable level of cooperation becomes wider. Thus, a positive-feedback structure ($g_{FB}^* > 0$) promotes cooperation by weakening the demand of Hamilton's rule for constitutive cooperation and allowing stable cooperation in a wider range of population structures, while a negative-feedback structure ($g_{FB}^* < 0$) makes this demand more stringent.

**Global invasion analysis shows that a positive-feedback structure improves the stability against public-good null mutants**

The local ESS analysis tests invasibility to small-effect mutations that change the quantitative features of the regulatory network. To understand the intuition behind the above results, we also tested invasion of any strategy in the plane of ($g_0 \times g_{FB}$) by public-good null mutants, i.e. ($g_0 = 0, g_{FB} = 0$). Again, for simplicity, we used the linear fitness function (Eq. (S14)) and assumed the same population structure. Recalculating Eqs. (S9),(S11),(S12), modified to the current scenario, we compared the fitness of an invading cheater in a resident cooperator patch to the fitness of a cooperator in a pure cooperator patch. We found that invasion of any strategy ($g_0, g_{FB}$) by the public-good null cheater ($g_0 = 0,\ g_{FB} = 0$) is independent of the basal public-good production rate $g_0$, and depends only on the feedback strength $g_{FB}$. The condition for stability of the strategy ($g_0, g_{FB}$) to invasion by the public-good null cheater ($g_0 = 0,\ g_{FB} = 0$) is that the relatedness should be higher than the exact same threshold obtained by the ESS analysis (Eq. (S19)), i.e., $r > r_{th}(g_{FB}) = \frac{1-g_{FB}}{\frac{B}{C}-g_{FB}}$ (this corresponds to Fig. 1C in the main text).

**Application to another example of intercellular feedbacks – quorum sensing regulation**

In this section, we analyze stability of two quorum sensing (QS) strategies; with and without a feedback on the QS signal, to invasion by response-null mutants and full QS-null mutants (response- and signal-null).

We first considered a simple QS design, where the QS signal is produced constitutively at a rate $y = y_0$ (both by QS cooperators and response-null mutants, but not by QS-null mutants where $y = 0$) and degraded at a rate $\gamma_S$. The ODE on the signal concentration $S$, and the steady state $S_{st}$ are therefore:

(S20) $$\frac{dS}{dt} = N\bar{y} - \gamma_S S,$$

(S21) $$S_{st} = \frac{N}{\gamma_S}\bar{y}.$$

We assumed that the public-good production is regulated by QS signals, and thus their production rate (of cooperators) is proportional to the signal concentration: $x = \tilde{\alpha} S_{st} = \alpha\bar{y}$ (where $\alpha = \tilde{\alpha}\frac{N}{\gamma_S}$). It is easy to show that the condition for stability of QS cooperators against full QS-null mutants is:

(S22) $$r > 1 - \sqrt{1 - \frac{C}{B}}.$$

This corresponds to a relatedness threshold that is lower than the threshold for stability of simple cooperators against cheaters which is $r_{th} = \frac{C}{B}$, see previous section and Eq. (S19)). Therefore, the simple QS design by itself provides an increased immunity against full QS-null mutants because QS cooperators can "sense" their presence and reduce cooperative effort accordingly. However, simple QS cooperators cannot "sense" the presence of response-null mutants that have a functional signaling gene (it is easy to show that the condition for stability against them is $r_{th} = \frac{C}{B}$). These mutants continue to produce QS signals and coerce cooperators to invest full effort in cooperation.

We therefore considered a feedback regulation in QS, such that the QS-signal production rate $y$ is regulated by the signal concentration $S$. By replacing $S$ instead of $PG$, $y$ instead of $x$, and $\gamma_S$ instead of $\gamma$ in Eqs. (S1)-(S4), we obtained the following relations:

(S23) $$y = f(\vec{\tilde{g}}, S) = \tilde{g}_0 + \tilde{g}_{FB} S,$$

(S24) $$\frac{dS}{dt} = \sum_{i=1}^{N} y_i - \gamma_S S,$$

(S25) $$S_{st} = \frac{N}{\gamma_S}\bar{y} = \frac{1}{\gamma_S}\sum_{i=1}^{N} y_i.$$

(S26) $$y = f(\vec{g}, \bar{y}) = g_0 + g_{FB}\bar{y} \qquad \left(g_0 = \tilde{g}_0 \ \ , \ \ g_{FB} = \tilde{g}_{FB}\frac{N}{\gamma_S}\right).$$

Again, we assumed that the public-good production rate is proportional to the steady state signal concentration $S_{st}$, with a proportionality constant $\alpha\tilde{g}_{FB}$ (assuming that as signaling molecules, public goods are also activated by signal-bound receptors, but that there is no basal public-good production rate):

(S27) $$x = \alpha\tilde{g}_{FB}S_{st} = \alpha g_{FB}\bar{y}.$$

The steady-state public-good production rate in a monomorphic population is thus:

(S28) $$x = \bar{x} = \alpha g_{FB}\bar{y} = \alpha g_{FB}\frac{g_0}{1-g_{FB}},$$

and in a dimorphic population:

(S29) $$x_i = \alpha g_{FB,i}\bar{y} = \alpha g_{FB,i}\frac{pg_{0,1}+(1-p)g_{0,2}}{1-pg_{FB,1}-(1-p)g_{FB,2}},$$

(S30) $$\bar{x} = px_1 + (1-p)x_2.$$

We proceeded to test the stability of QS feedback strategies $\left(g_{0,2} \equiv g_0\ ,\ g_{FB,2} \equiv g_{FB}\right)$ to invasion by response-null mutants with a functional signaling gene. These mutants continue to produce the basal level of QS signals ($g_{0,1} = g_0$) but lack the receptor and therefore do not produce further signals ($g_{FB,1} = 0$), and do not produce public goods at all ($x_1 = 0$). The resulting condition for stability of QS feedback strategies against these response-null mutants is identical to the simpler case of auto-regulated public goods (i.e., $r > \frac{1-g_{FB}}{\frac{B}{C}-g_{FB}}$, see previous section and Eq. (S19)). The stabilizing effect of the intercellular positive-feedback structure applies here as well, improving the stability against response-null mutants that are problematic to the simple QS design.

### Supplementary tables

**Table S1 – *B. subtilis* strain list**

| Strain name | Genotype | Source reference |
|---|---|---|
| AES101 | *B. subtilis* PY79 wild-type | Bacillus genetic stock center |
| BD1916 (*B. subtilis* 168) | $P_{spac}$-srfA | Kind gift from D. Dubnau [7] |
| AES827 | zjd-89::(Tn917::pTV21Δ2::pD177.1::pD179.1 Kan Cm) | [8] |
| AES2026 | amyE::($P_{srfA}$-3xYFP Sp) | [9] |
| AES2030 | sfp::($sfp_{3610}$ Sp) swrA::$swrA_{3610}$ lacA::($P_{43}$-YFP MLS) | [9] |
| AES2135 | ΔcomQXP::Tet | [9] |
| AES2137 | sfp::($sfp_{3610}$ Sp) swrA::$swrA_{3610}$ lacA::($P_{43}$-YFP MLS) ppsB::($P_{trpE}$-mCherry Ph) | [9] |
| AES2663 | ppsB::($P_{trpE}$-mCherry Ph) | [9] |
| AES2864 | zjd-89::Tn917::pTV21Δ2::pD177.1::pD179.1 Kan Cm) ΔcomQXP::Tet | AES2135→AES827 |
| AES2965 | zjd-89::($P_{hs}$-comXP Kan Cm Sp) ΔcomQXP::Tet | AEC1054→AES2864 |
| AES2970 | zjd-89::($P_{srfA}$-comXP Kan Cm Sp), ΔcomQXP::Tet | AEC1057→AES2864 |
| AES3354 | ΔcomXPA::Tet | This study – (LFH PCR) |
| AES3380 | ΔcomXP::Tet | This study – (LFH PCR) |
| AES3412 | ΔcomXPA::Tet amyE::($P_{srfA}$-3xYFP Sp) | AES2026→AES3354 |
| AES3414 | ΔcomXP::Tet amyE::($P_{srfA}$-3xYFP Sp) | AES2026→AES3380 |
| AES3425 | ΔcomXPA::Tet amyE::($P_{srfA}$-3xYFP Sp) zjd-89::($P_{hs}$-comXP Kan Cm Sp) | AES2965→AES3412 |
| AES3427 | ΔcomXPA::Tet amyE::($P_{srfA}$-3xYFP Sp) zjd-89::($P_{srfA}$-comXP Kan Cm Sp) | AES2970→AES3412 |
| AES3429 | ΔcomXP::Tet amyE::($P_{srfA}$-3xYFP Sp) zjd-89::($P_{hs}$-comXP Kan Cm Sp) | AES2965→AES3414 |
| AES3430 | ΔcomXP::Tet amyE::($P_{srfA}$-3xYFP Sp) zjd-89::($P_{srfA}$-comXP Kan Cm Sp) | AES2970→AES3414 |
| AES3439 | ΔcomXP::Tet amyE::($P_{srfA}$-3xYFP Sp) zjd-89::($P_{hs}$-comXP Kan Cm Sp) ppsB::($P_{trpE}$-mCherry Ph) | AES2663→AES3429 |

| | | |
|---|---|---|
| AES3440 | ΔcomXP::Tet amyE::($P_{srfA}$-3xYFP Sp) zjd-89::($P_{srfA}$-comXP Kan Cm Sp) ppsB::($P_{trpE}$-mCherry Ph) | AES2663→AES3430 |
| AES3628 | sfp::($sfp_{3610}$ Sp) swrA::$swrA_{3610}$ lacA::($P_{43}$-YFP MLS) ΔcomXP::Tet | AES3380→AES2030 |
| AES3634 | sfp::($sfp_{3610}$ Sp) swrA::$swrA_{3610}$ lacA::($P_{43}$-YFP MLS) ΔcomXP::Tet zjd-89::($P_{srfA}$-comXP Kan Cm Sp) | AES2970→AES3628 |
| AES3927 | zjd-89::($P_{spac}$-comXP Kan Cm Sp) ΔcomQXP::Tet | AEC1174→AES2864 |
| AES3930 | sfp::($sfp_{3610}$ Sp) swrA::$swrA_{3610}$ ppsB::($P_{trpE}$-mCherry Ph) lacA::($P_{43}$-YFP MLS) ΔcomXPA::Tet | AES3354→AES2137 |
| AES3938 | sfp::($sfp_{3610}$ Sp) swrA::$swrA_{3610}$ lacA::($P_{43}$-YFP MLS) ΔcomXP::Tet zjd-89::($P_{spac}$-comXP Kan Cm Sp) | AES3927→AES3628 |
| AES3940 | sfp::($sfp_{3610}$ Sp) swrA::$swrA_{3610}$ ppsB::($P_{trpE}$-mCherry Ph) lacA::($P_{43}$-YFP MLS) ΔcomXPA::Tet zjd-89::($P_{spac}$-comXP Kan Cm Sp) | AES3927→AES3930 |
| AES3944 | sfp::($sfp_{3610}$ Sp) swrA::$swrA_{3610}$ ppsB::($P_{trpE}$-mCherry Ph) lacA::($P_{43}$-YFP MLS) ΔcomXPA::Tet zjd-89::($P_{srfA}$-comXP Kan Cm Sp) | AES2970→AES3930 |

**Table S2** – ***E. coli* strain list**

| Strain name | Genotype | Source reference |
|---|---|---|
| AEC777 | pDR111 | [10] |
| AEC945 | pDL30::$P_{srfA}$-3xYFP | [11] |
| AEC1054 | pDR111::$P_{hs}$-comXP | This study |
| AEC1057 | pDR111::$P_{srfA}$-comXP | This study |
| AEC1174 | pDR111::$P_{spac}$-comXP | This study |

**Table S3** – **Primer list**

| | |
|---|---|
| comXP-del-P1 | TGAAGGAGATTGTGGAGCAA |
| comXP-del-P2 | TGCCCGCAGCTGTGACAATGTTTCAGTTTCTTGCACATCA |
| comXP-del-P3 | GTAGCGCGGTGGTCCCACGGCTTTAGATGGGCGCCT |
| comXP-del-P4 | GGTTGGCGTTAATCTCCAAACCAAC |
| comXPA-del-P1 | TGAAGGAGATTGTGGAGCAA |
| comXPA-del-P2 | TGCCCGCAGCTGTGACAATGTTTCAGTTTCTTGCACATCA |
| comXPA-del-P3 | GTAGCGCGGTGGTCCCACAGCGGTCCATTGAATACAGC |
| comXPA-del-P4 | GGTGAGCCGGTGATGTTTAC |
| comXP-F | AGCTGCTAGCAAAGGGGGATACAAGATGC |
| comXP-R | CATGGCATGCTGTTTTCCTCCCTTTTACTCAC |
| Pspac-F | ATCTGAATTCTACACAGCCCAGTC |
| Pspac-R | ACTAGCTAGCCCGAAATAGCCCAAAGTA |

### Supplementary references


1. Dieckmann U (1997) Can adaptive dynamics invade? *Trends Ecol Evol* **12**: 128–131.

2. Maynard Smith J (1982) *Evolution and the theory of games Cambridge*. Cambridge University Press.

3. Smith J, Van Dyken JD, Zee PC (2010) A generalization of hamilton's rule for the evolution of microbial cooperation. *Science (80- )* **328**: 1700–1703.

4. Brown SP (1999) Cooperation and conflict in host-manipulating parasites. *Proc R Soc B Biol Sci* **266**: 1899–1904.

5. Brown SP, Johnstone RA (2001) Cooperation in the dark: signalling and collective action in quorum-sensing bacteria. *Proc R Soc B Biol Sci* **268**: 961–965.

6. West SA, Griffin AS, Gardner A, Diggle SP (2006) Social evolution theory for microorganisms. *Nat Rev Microbiol* **4**: 597–607.

7. Hahn J, Dubnau D (1991) Growth stage signal transduction and the requirements for srfA induction in development of competence. *J Bacteriol* **173**: 7275–7282.

8. Vin YY, Kondratiev DE (2012) Information approach to studying byzantine law and its receptions. *Byzantinoslavica* **70**: 76–96.

9. Even-Tov E, Omer Bendori S, Valastyan J, Ke X, Pollak S, Bareia T, Ben-Zion I, Bassler BL, Eldar A (2016) Social Evolution Selects for Redundancy in Bacterial Quorum Sensing. *PLoS Biol* **14**:.

10. Britton RA, Eichenberger P, Gonzalez-Pastor JE, Fawcett P, Monson R, Losick R, Grossman AD (2002) Genome-wide analysis of the stationary-phase sigma factor (Sigma-H) regulon of Bacillus subtilis. *J Bacteriol* **184**: 4881–4890.

11. Omer Bendori S, Pollak S, Hizi D, Eldar A (2015) The RapP-PhrP quorum-sensing system of Bacillus subtilis strain NCIB3610 affects biofilm formation through multiple targets, due to an atypical signal-insensitive allele of RapP. *J Bacteriol* **197**: 592–602.

### Supplementary figures

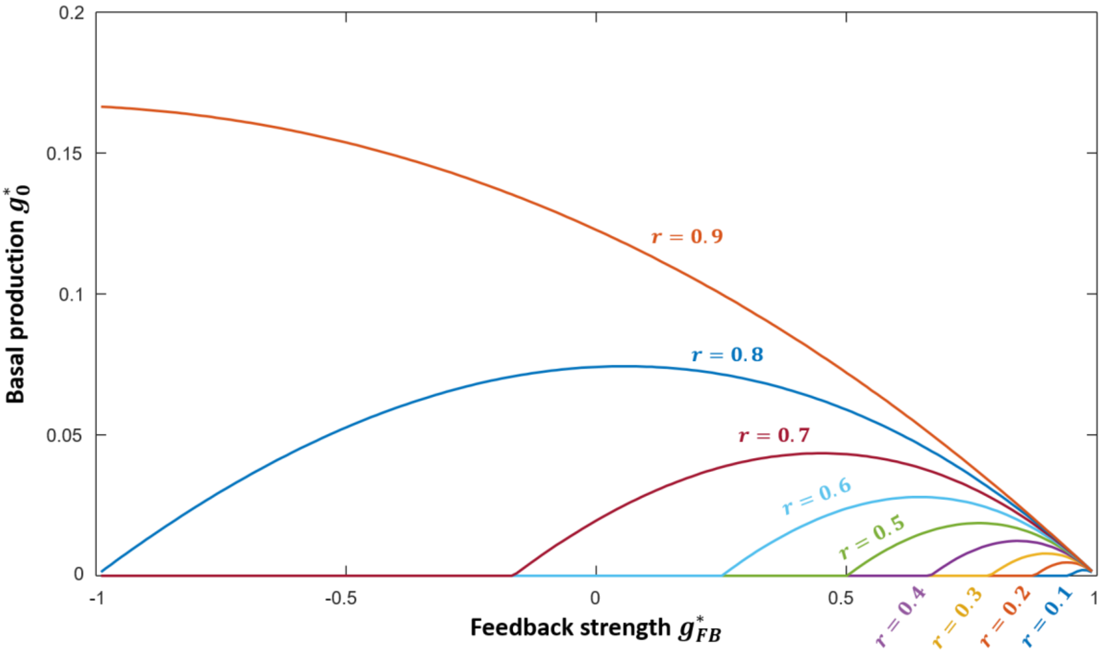


**Figure S1 – Locally evolutionarily-stable regulatory strategies of public-good production.** Shown are curves of local ESSs (for varying relatedness levels), in the plane of feedback strategies with a basal public-good production rate $g_0^*$ and feedback strength $g_{FB}^*$. Going left from $g_{FB}^* = 1$, each curve meets the X-axis at some point and then goes along the axis, corresponding to evolutionarily stable cheating (for $r = 0.9$ this happens at a point smaller than $g_{FB}^* = -0.1$). The benefit to cost ratio parameter used is $\frac{B}{C} = 1.5$.

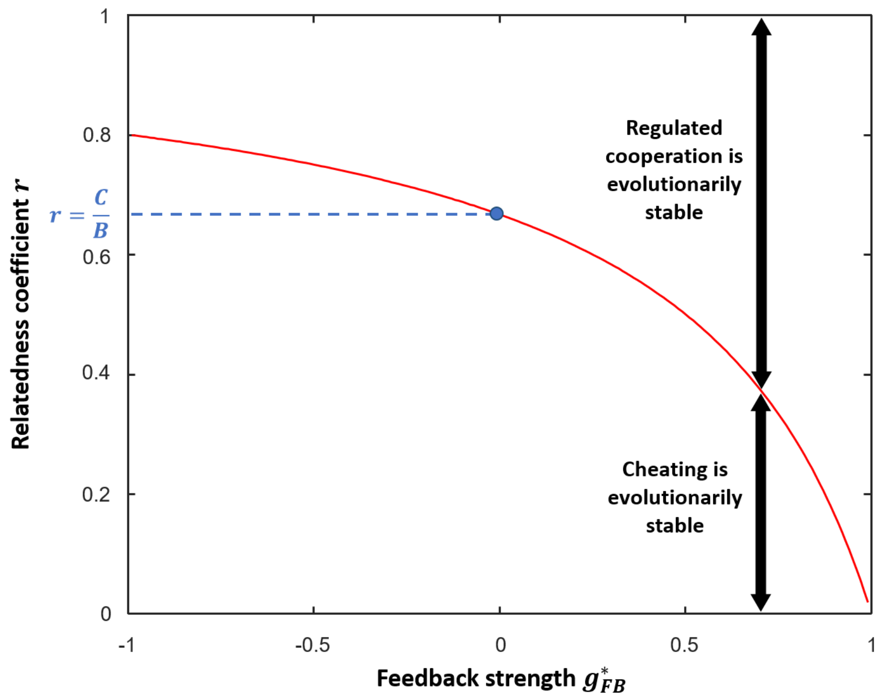


**Figure S2 – Threshold relatedness as a function of feedback slope $r_{th}(g^*_{FB})$ for having locally evolutionarily-stable cooperation.** For every $g^*_{FB}$ there is a positive evolutionarily-stable level of cooperation ($x^* > 0$) only when the relatedness in the population is higher than the threshold $r_{th} = \frac{1-g^*_{FB}}{\frac{B}{C}-g^*_{FB}}$ (red line), otherwise there will be a zero-cooperation ESS ($x^* = 0$). The blue dot represents the Hamilton rule threshold for constitutive cooperation $r_{th}(g^*_{FB} = 0) = \frac{C}{B}$. The bebefit to cost ratio paramter used here is $\frac{B}{C} = 1.5$.

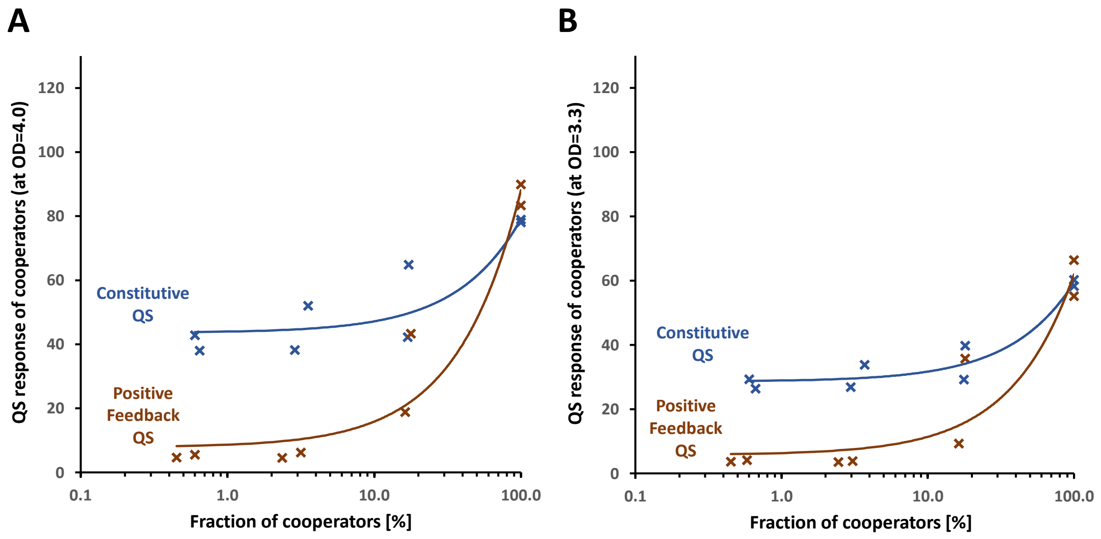


**Figure S3 – Co-culture gene-expression trends are independent of the interpolating bacterial density.** Similar to Fig. 2E in the main text but for different choices of interpolating bacterial density, equivalent to $OD_{600}$ = 4.0 in **(A)** and $OD_{600}$ = 3.3 in **(B)**.

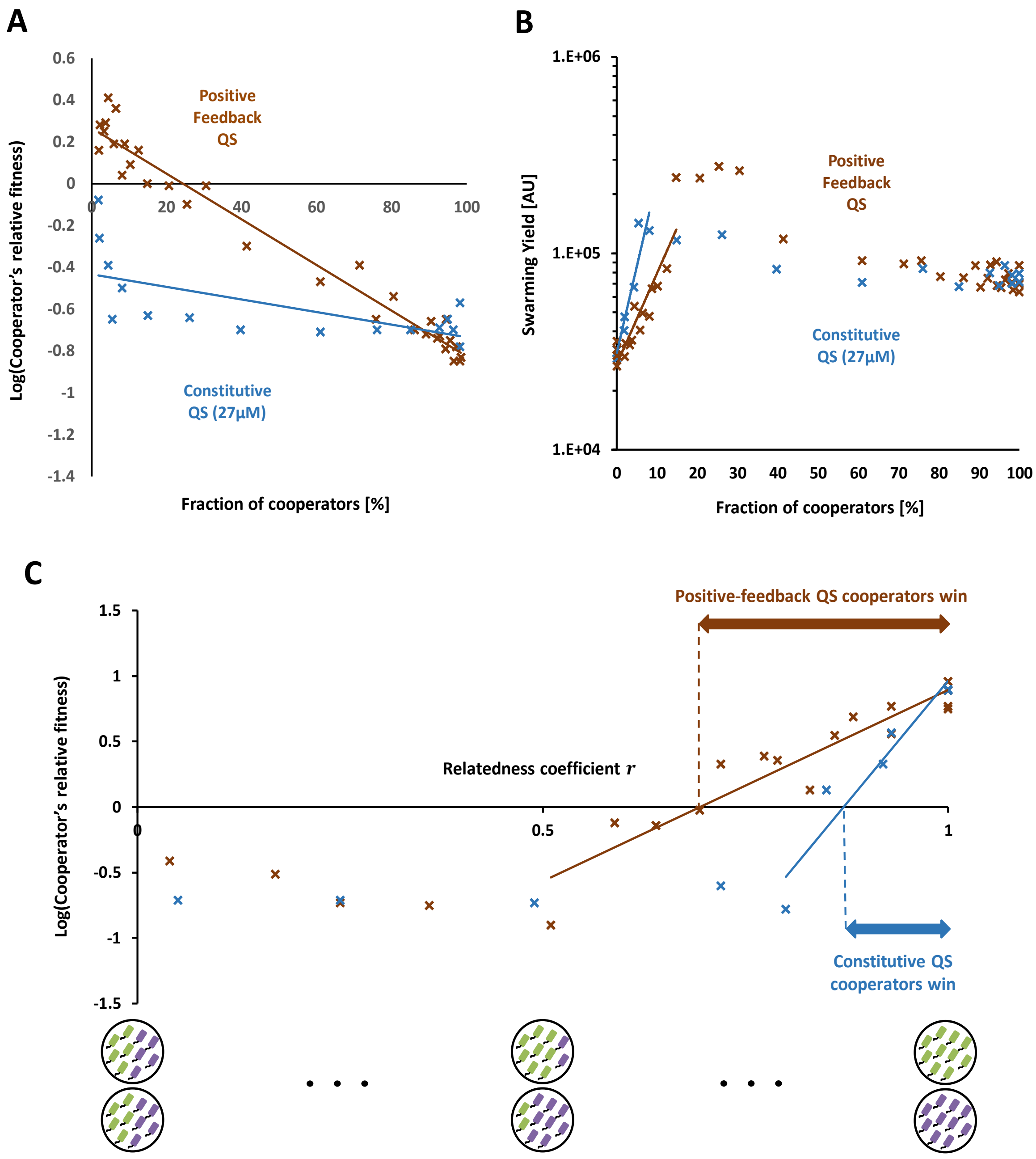


**Figure S4 – Competition results are weakly dependent on IPTG concentration.** Similar to Fig. 3 in the main text but with 27μM IPTG for the constitutive-QS cooperator (two biological repeats) (data of positive-feedback-QS cooperator is presented again for comparison). For the added constitutive-QS cooperator data, $R^2 = 0.42$ in **(A)**, $R^2 = 0.90$ in **(B)**, and $R^2 = 0.91$ in **(C)**.
